# Scalable large-area mesh-structured microfluidic gradient generator for drug testing applications

**DOI:** 10.1101/2022.07.14.500002

**Authors:** Shital Yadav, Pratik Tawade, Ketaki Bachal, Makrand A. Rakshe, Yash Pundlik, Prasanna S. Gandhi, Abhijit Majumder

## Abstract

Microfluidic concentration gradient generators are useful in drug testing. drug screening, and other cellular applications to avoid manual errors, save time, and labor. However, expensive fabrication techniques make such devices prohibitively costly. Here, in the present work, we developed a microfluidic concentration gradient generator (μCGG) using a recently proposed non-conventional photolithography-less method. In this method, ceramic suspension fluid was shaped into a square mesh by controlling Saffman Taylor instability in a Multiport Lifted Hele-Shaw Cell (MLHSC). Using the shaped ceramic structure as template, the microfluidic concentration gradient generator (μCGG) was prepared by soft lithography. The concentration gradient was characterized and effect of the flow rates were studied usingCOMSOL simulations. The simulation result was further validated by creating fluorescein dye (Fluorescein isothiocanate, FITC) gradient in the fabricated μCGG. To demonstrate the use of this device for drug testing, we created various concentrations of an anticancer drug - curcumin - using the device and determined its inhibitory concentration on cervical cancer cell-line HeLa. We found that the IC50 of curcumin for HeLa to be 28.6 ± 6.1 μM which matched well with the conventional muti-well drug testing method (34.9 ± 1.7 μM). This method of μCGG fabrication has multiple advantages over conventional photolithography such as: i) the channel layout and inlet-outlet arrangements can be changed by simply wiping the ceramic fluid before it solidifies, (ii) it is cost effective, (iii) large area patterning is easily achievable, and (iv) the method is scalable. This technique can be utilised to achieve broad range of concentration gradient to be used for various biological and non-biological applications.

**Table of Content:** 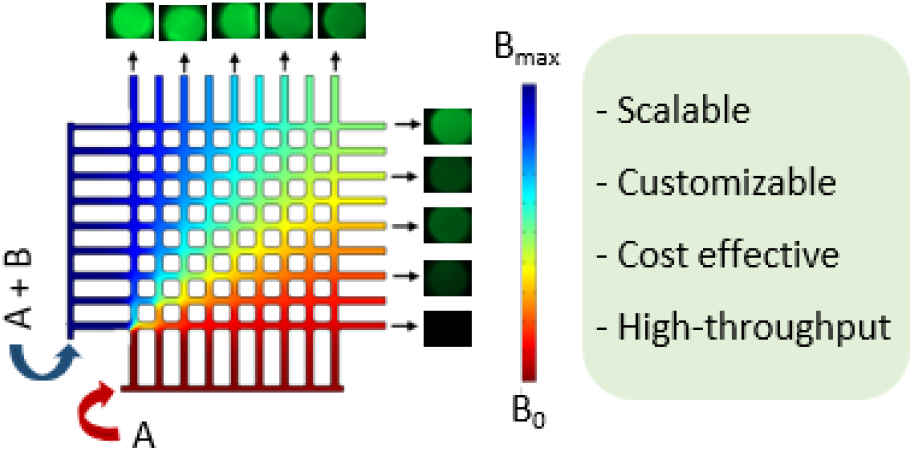

## 1. Introduction

Microfluidics concentration gradient generators (μCGG) are widely used for various chemical and biological studies including chemical synthesis ^1^, serial dilution ^2,3^, sensors ^4,5^, clinical diagnostics ^6^, chemotaxis ^7^, drug screening ^8^, biochemical assays, etc. μCGG provides precise control over concentration of the active molecule in space and time ^9^. In such flow-based systems, mostly two fluid streams are combined and allowed to mix or diffuse, in ‘T’ or ‘Y’ shaped channels ^10^. Christmas tree design is another widely used flow-based device where gradient is generated by continuous splitting and recombining fluid streams ^10^. The well-controlled shapes of μCGG are commonly fabricated using lithography-based techniques, including photolithography, soft lithography, e-beam lithography, etc. These methods comprise of multi-step process and need sophisticated instruments, thereby increasing processing cost and limits industrial scale-up ^11^. Recently, in order to overcome these limitations, nonconventional methods have also been reported including paper-based ^12,13^ or thread-based ^14^ gradient generation. However, in both paper and thread-based μCGG, it is difficult to collect the outlet stream as fluid is absorbed in the device.

In this work, we report the fabrication, optimization, and validation of a flow-based μCGG for drug screening application. The device was made with poly-dimethyl siloxane (PDMS) based soft-lithography technique using a template which was fabricated using a non-conventional method of shaping ceramic fluid in a lifted Hele-Shaw cell as reported earlier ^15^. Controlled shaping of fluid allows customising number of outlets, customising shape by wiping fluid before crosslinking and area of pattern, thereby controlling gradient strength in μCGG. The optimum operating conditions to generate a concentration gradient were determined using COSMOL simulations and validated by experiment with fluorescent dye FITC. The outlet concentration profile was shown to be controlled by changing the flow rate ratios of both inlets. Further, the inhibitory concentration (IC50) of curcumin on cervical cancer cells (HeLa) was estimated using the fabricated μCGG (28.6 ± 6.1 μM). The result matched well with the same estimated using conventional drug testing platforms i.e., multi-well plates (34.9 ± 1.7 μM). We propose that this easy-to-fabricate and scalable μCGG can be used for the generation of concentration gradients for various biological and non-biological applications.

## 2. Material and methods

### 2.1 Fabrication of square mesh microfluidic device

Fabrication is carried out in a parallel lifting Hele Shaw cell custom-built at Suman Mashruwala Advanced Microengineering Laboratory, IIT Bombay. The details of the setup can be found in ^15^. The cell plates of the setup are modified to suit the proposed application.

#### 2.1.1. Controlling source holes

Using a CNC-micro drilling machine source-holes were drilled on one of the cell plates for fabricating the proposed μCGG structures. Diameter of source-holes in the present study is kept equal to 0.5 mm for all the experiments. Locations of these holes were based on the guidelines given in ^16^ such that we would evolve a square mesh structure after the process is done. The square mesh with several sizes and pitch dimensions are prepared.

#### 2.2.2 Square mesh preparation

The measured amount of shear thinning suspension consisting of ceramic (Alumina) particles suspended in photoresist (HDDA) [as reported in ^16^] (fig 1.a.i), is squeezed between two cell plates to the required thickness and radius (fig 1.a.ii). The array of source holes on one of the cell plates is decided as mentioned in 2.2.1. During the squeezing step, all the source holes are properly sealed. Squeezing is followed by a small delay to neutralise the normal stresses within the fluid. The cell plates are separated with unsealed holes (fig 1.a.iii). As the two plates start separating, air moves inside from the source holes, creating the air finger. Each air finger growth is shielded by adjacent air finger which finally gives square mesh pattern due to pre-designed source hole locations ^17^. The Saffman-Taylor instability (viscous fingering) phenomenon is controlled by conducting fabrication with a proper capillary number and aspect ratio ^15^. Once the plate separation is completed, we get a square mesh pattern on each plate. Channels for inlet and outlet streams are created by wiping selected areas. Presence of photo-initiator in the ceramic suspension allows it to UV cure (fig 1.a.iv) and finally we get cross-linked square mesh pattern. Figure 1.a.v shows the digital image of a 9X9 square mesh template used as the template for the study. Changing distance between drilled holes and number of holes can be utilised to customise the pattern and patterned area.

**Figure 1:**
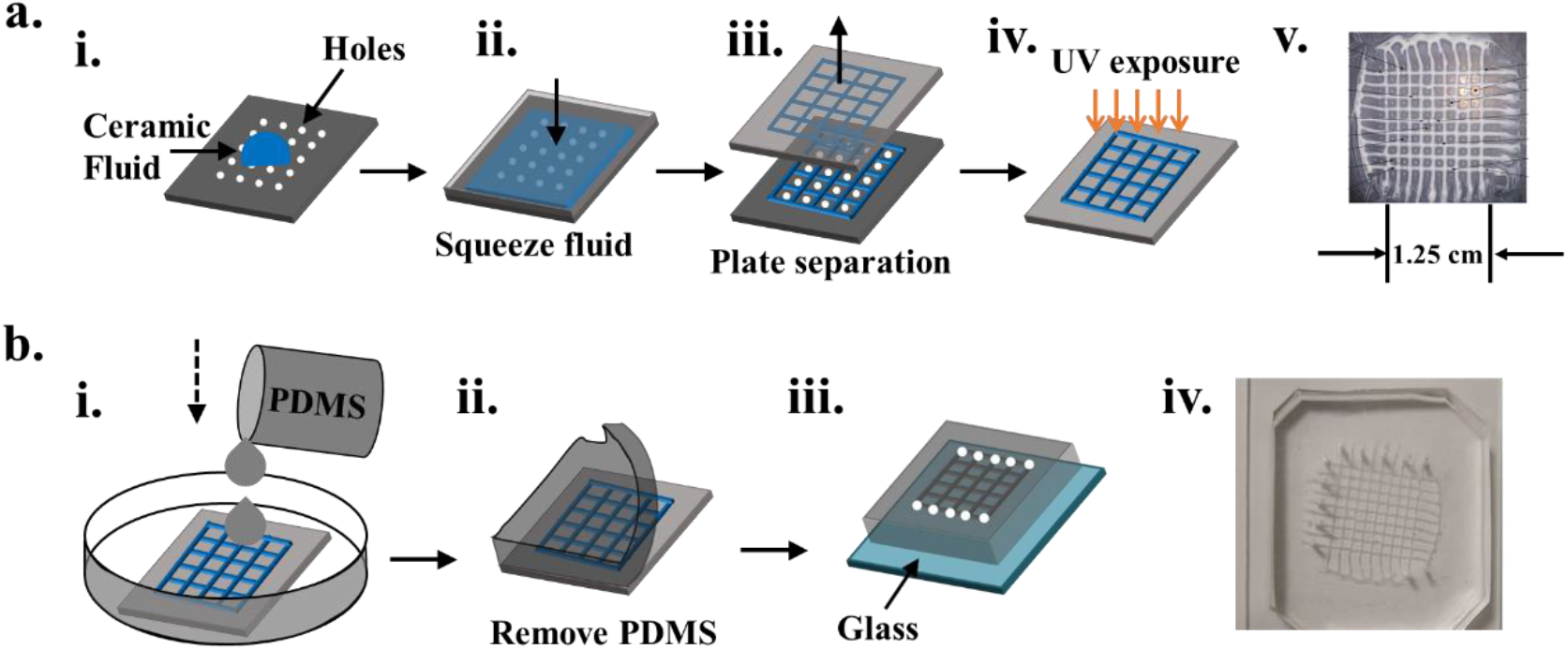
Schematic of preparation of (a) template and (b) microfluidic device

#### 2.1.3 Preparation of microfluidic device

PDMS (Sylgard 184 from Dow Corning) is mixed in 10:1 (w/w) with crosslinker and then degassed under vacuum. This uncured PDMS was then poured onto square mesh pattern prepared (fig 1.b.i) and cured at 60 °C for 12hrs. The casted cured PDMS is then peeled off from template (fig 1.b.ii) and is referred as negative replica. In order to make channels, this negative replica is plasma oxidised and bonded glass slide by bringing two surfaces in conformal contact. For flow, inlets and outlets were made using biopsy punches as shown in fig 1.b.iii, before bonding with the glass. Figure 1.b.iv shows photo of fabricated microfluidic device for CGG.

### 2.2 Template characterisation

The height and width of the channels and nodes were measured using white light interferometry (WLI). WLI results in 3D profile of area scanned.

### 2. 3. Flow setup

The square mesh-shaped microfluidic device can be customised in terms of inlet-outlet number and position. In this work, fabricated μCGG have two inlet ports and 10 outlet ports – alternate channels (fig 2.a). Capillary tubes of 1mm diameter were used for connecting the syringe and the inlet ports of the device. Outlets were collected in 200 μl microtips (fig 2.b). Syringe pump (Era’s NE-1000) was used for the experiment, programmed for the desired flow rate.

**Figure 2:**
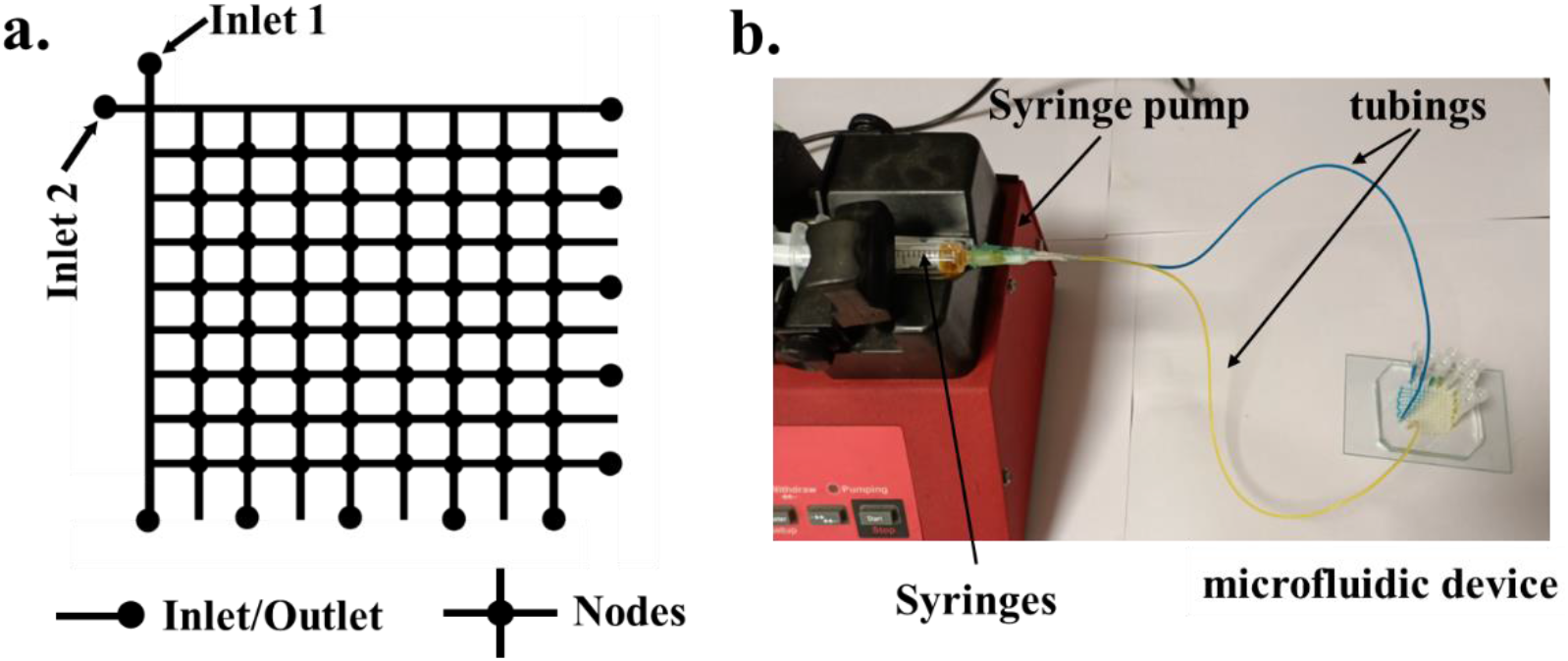
Schematic of (a) inlet-outlet positions and (b) experimental setup for μCGG

### 2.4 Characterisation of concentration gradient

#### 2.4.1. COMSOL simulation

The 2D geometry of the device is made in COMSOL Multiphysics geometry module. The meshing is done to discretize the domains into small elements, as is done in finite element method. The concentration at the outlets was simulated using COMSOL (version 5.2) assuming laminar flow conditions. Other COMSOL parameters are mentioned in Table 1. In addition to equal flow rates in both inlets, we also checked the effect of the inlet flow rate ratio on the outlet concentration profile (Table 1).

**Table 1:** COMSOL simulation parameters

|  |  |
| --- | --- |
| Software | COMSOL Multiphysics 4.4 |
| Physics type (In COMSOL) | Laminar flow & Transport of diluted species |
| **Material** | Water |
| Viscosity | 0.001 Pa s |
| Density | 1000 kg m<sup>-3</sup> |
| **Material** | FITC |
| FITC diffusivity in water | 4.25X 10<sup>-10</sup> m<sup>2</sup>/s |
| **Boundary conditions** |  |
| Inlet velocity condition (equal inlet velocity) | 0.001 – 0.02 m/s |
| Inlet1:Inlet2 flow rate | 50-100, 50-150, 50-200, 200-50, 150-50, 100-500 μl/min |
| Inlet concentration | 0 and 0.05 mol/m<sup>3</sup> |
| Outlet condition | Constant pressure |
| Walls | No slip condition |

#### 2.4.2. Fluorescein dye gradient

Sodium fluorescein dye (FITC-376 Da) in MilliQ water was used to generate gradients at different flow rates in the fabricated microfluidic device. The collected outlet is imaged in a fluorescence microscope (EVOS FL-Auto). The concentration was determined by comparing fluorescence intensities of known concentration i.e., standard curve of intensity vs concentration, (supplementary figure S1) with intensities from outlets from device.

#### 2.4.3. Curcumin gradient

The concentration gradient of model drug-Curcumin (389 Da), was generated in cell growth media at flow rate 50μl/min. The absorbance of the outlet solution was measured at 425 nm using a SpectraMax M2^e^ microplate reader. To estimate outlet concentration, standard curve with absorbance or known concentration was plotted (supplementary figure S2) and outlet concentration of curcumin from device was determined.

### 2.5 Cell culture

The cervical cancer cells (HeLa) were cultured in High glucose Dulbecco’s Modified Eagle Medium (DMEM) supplemented with 1% Anti-Anti, 1% L-glutamine and 20% Fetal Bovine Serum (Himedia). Cells are trypsinized with 0.05 % trypsin - EDTA(1X) (Gibco), incubated for 5 mins at 37 °C. To get pellet of cells it was centrifuged (1000rpm, 5min). After centrifugation, cells are resuspended with fresh medium and counted using haemocytometer and seeded in 96 well plate.

### 2.6 MTT assay

HeLa cells were seeded in 96 well cell culture plate at 5000 cells per well. After 24 h, cells were exposed to different concentrations of curcumin drug (in growth media) prepared by (i) μCGG and (ii) manual dilution. Cells were allowed to proliferate for 24 h and cell viability was estimated by quantifying reduction of tetrazolium dye – MTT (3-(4,5-dimethylthiazol-2-yl)-2,5-diphenyltetrazolium bromide) by monitoring absorbance at 590 nm using a SpectraMax M2^e^ Microplate reader. IC_50_ value i.e., 50% inhibition of cell viability – indicate cytotoxicity of drug on cells, was determined by nonlinear regression fitting of the concentration-dependent proliferation inhibitory data ^18^.

## 3. Results

### 3.1 Square mesh dimension

The dimensions of square mesh can be varied by changing the initial thickness of the fluid film and the pitch distance of the source holes in the plate ^15^. The optimised parameters for the preparation of square mesh are selected from source hole characterization graph ^15,16^. The intersections of branches in the mesh is referred to as nodes. The shape of the channels and the nodes is as shown in figure 3.a & b respectively. The template used in this study has an average channel height of 58.9 ± 5.75 μm and width of 650.7 ± 38.14 μm. The average height of node is 109.2 ± 22 μm as measured by white light interferometer (WLI).

**Figure 3:**
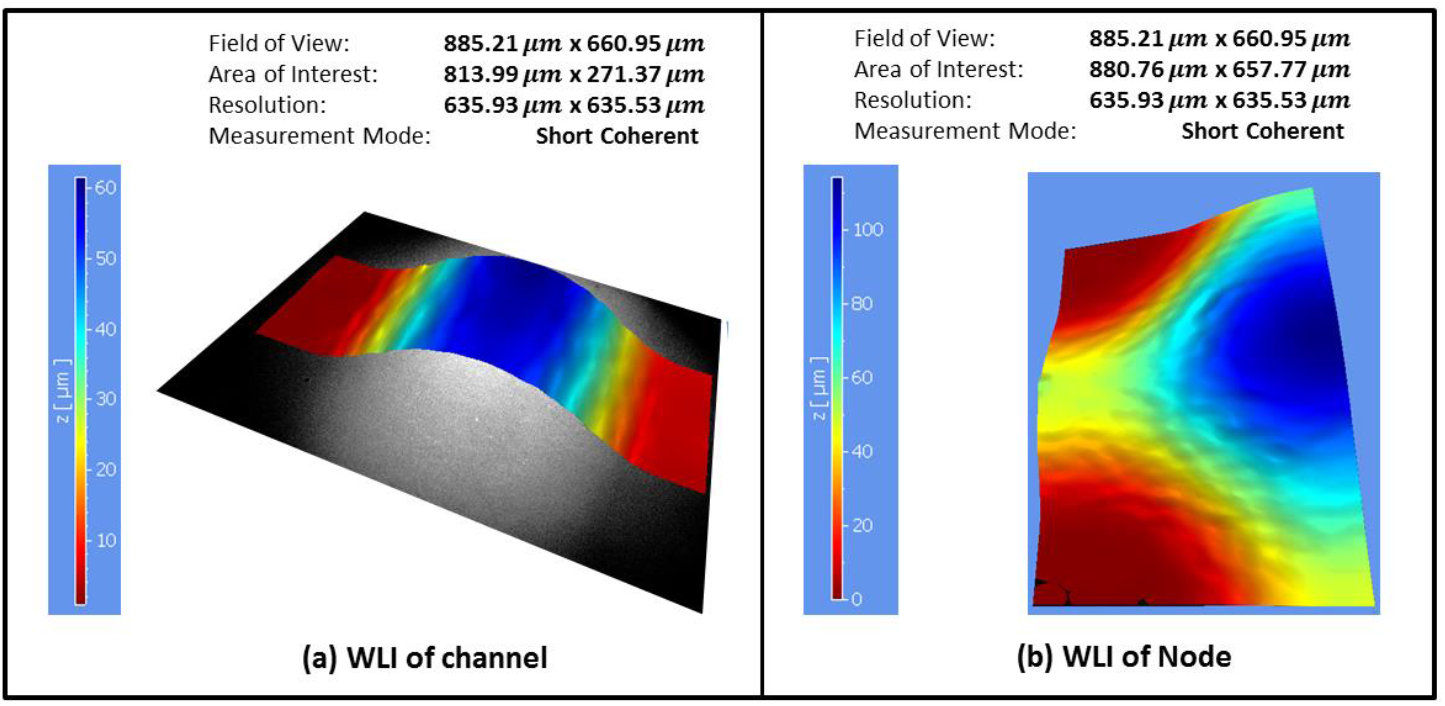
WLI image of the (a) channel and (b) node

### 3.2 Concentration gradient characterisation

Using COMSOL simulation, we analysed the concentration at the outlet streams at various flow rates. Figure 4.a shows the simulation result for the concentration gradient formed in the device with equal inlet flow rates i.e., 50 μl/min. The simulation result was verified with further experiments using 50 μM FITC (fluorescein dye) in water as one inlet and pure water as other, with an equal flow rate of 50 μl/min (fig 4.b) and 100 μl/min (fig 4.c) in both the inlets. Furthermore, the flow rate of one inlet was varied keeping the other constant (fig 4.d-e). Figure 4.f compiles different outlet concentration profiles obtained with different inlet flow rate ratios showing that the concentration profile can be controlled by changing inlet flowrates. Figure 4.g shows fluorescence variation across the device outlets for equal flow rate of 50 μl/min.

**Figure 4:**
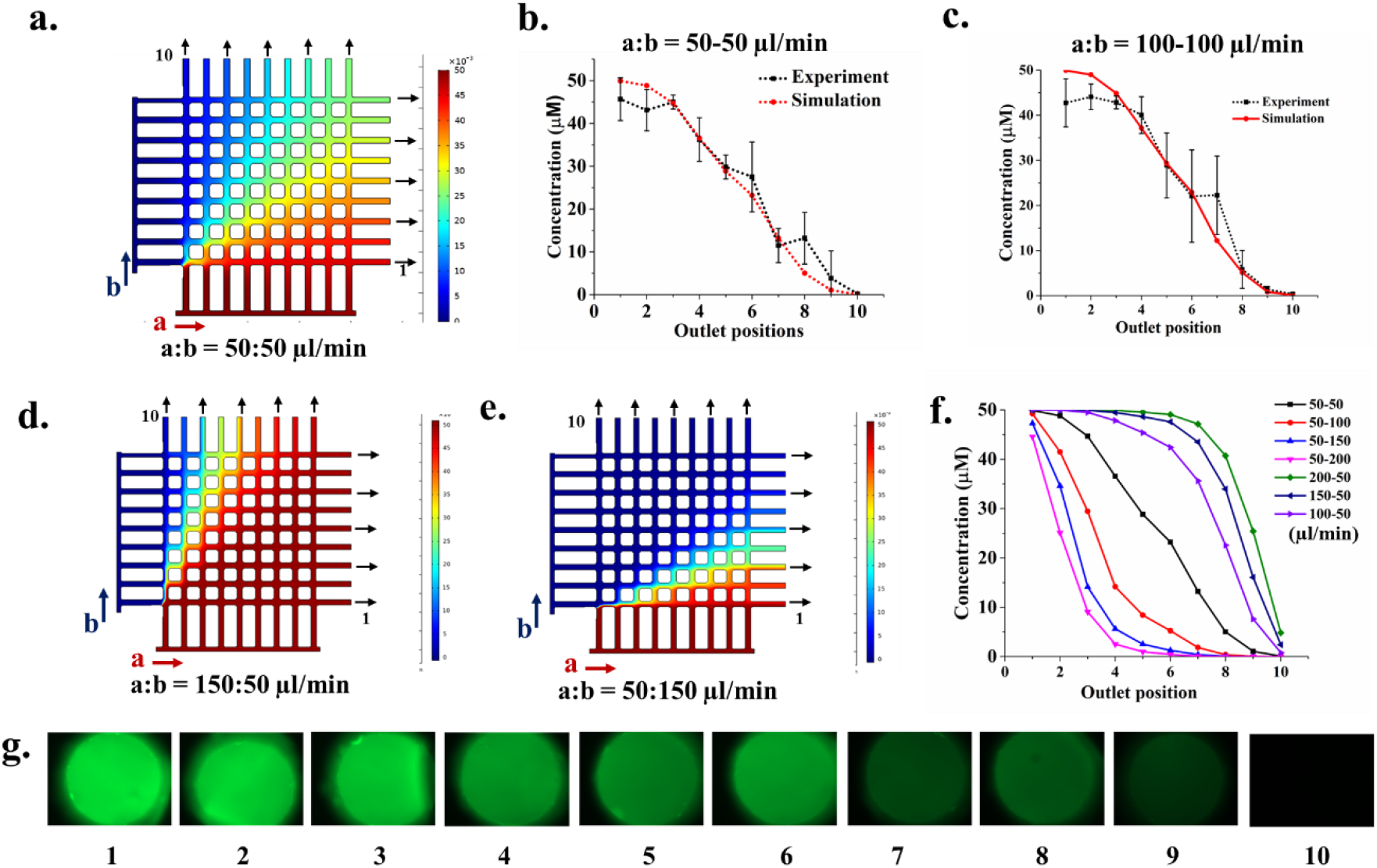
Gradient characterisation from COMSOL for (a) inlet *a:* 50 μl/min, inlet *b:* 50 μl/min i.e., equal inlet flow rate of 50μl/min; comparison between outlet concentrations obtained from experiment and simulation at flow rate of (b) inlet *a:* 50 μl/min, inlet *b*: 50 μl/min and (c) inlet *a:* 100 μl/min, inlet *b*: 100 μl/min for both the inlets. If both inlets have different flow rate, concentration profile changes as in (d) inlet *a:* 150 μl/min, inlet *b*: 50 μl/min and (e) inlet *a:* 50 μl/min, inlet *b*: 150 μl/min; (f) shows outlet concentrations with different combinations of inlet flow rates and (g) fluorescence intensity (green) of FITC gradient from experiment at inlet *a*: 50 μl/min, inlet *b*: 50 μl/min flow rate. (In COMSOL, red = 50 μM concentration of FITC, Blue = 0 μM concentration of FITC in water)

### 3.4. Drug gradient and its effect on cell metabolic activity

The potential application of concentration gradient is in determining the effect of various drugs/biomolecules on cells. We selected curcumin as model drug to study its cytotoxic effect on HeLa cells, a commonly used model cell line for cervical cancer. First, the concentration gradient of curcumin in growth media was generated in the fabricated device. The concentration of curcumin at each outlet was determined using absorbance of solution at 425nm (fig 5.a). Using MTT assay we then tested the cytotoxicity of curcumin on HeLa cells by measuring the percent inhibition w.r.to drug collected from the device outlets (fig. 5b). As the graph shows, drug concentration coming from outlet 5 gives the IC_50_ i.e., inhibitory concentration at which 50% cell are viable/dead. Comparing with figure 5.a, the concentration of solution from outlet number 5 is found to be ~32 μM. Therefore, IC_50_ of curcumin on HeLa cells is ~32 μM. To note, our COMSOL simulation estimated the concentration at outlet 5 is 28 μM which matches well with the experimental measurement (fig 4.c). Hence, drug concentration at different outlets can be estimated using COMSOL simulation by providing molecular weight and diffusivity of drug as input and thus eliminating the need of experimental measurement of drug concentration every time. Furthermore, to validate the device, IC_50_ of curcumin on Hela was estimated using a conventional 96 well plate in which concentration gradient was generated manually by serial dilution. - Using this conventional method, IC_50_ was found to be ~35 μM (figure 5.c) which matched well with IC50 estimated from our μCGG device (32 μM). This result indicates that the μCGG can potentially be used for concentration generation in drug testing and drug screening replacing the time consuming, labour intensive, and error-prone method of creating concentration gradients manually.

**Figure 5:**
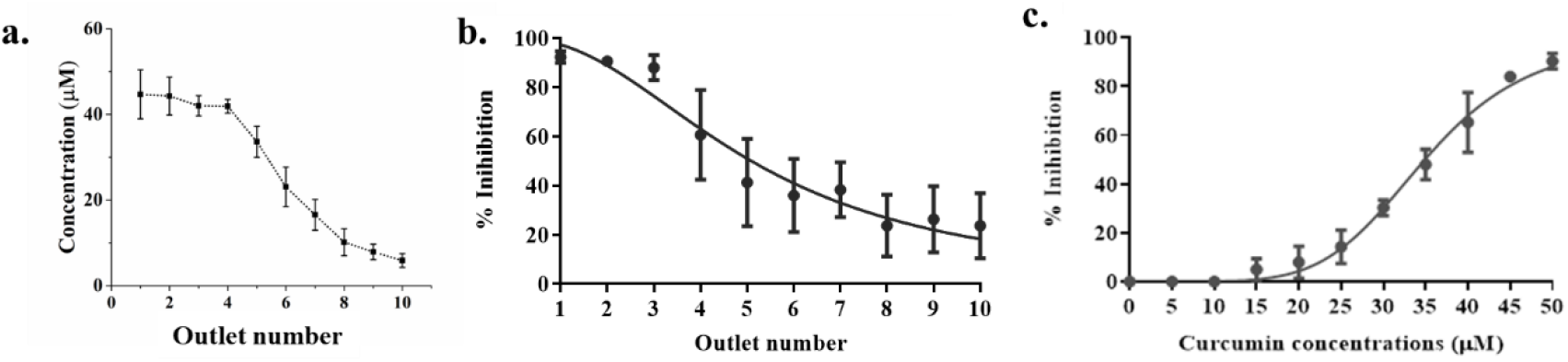
(a) Curcumin concentration at different outlets; Determination of IC50 using (b) fabricated μCGG and (c) conventional 96-well plate

### 3.5. Scalability of the Fabrication Process

Fabrication process used for the mesh pattern generation is spontaneous and scalable. As the shaping of fluid is created by varying air entrance pathways, changing the initial experimental parameter like initial fluid volume, initial fluid film thickness, initial squeezed fluid film radius and pitch distance of the array of source holes, can change the micro-meso scale structure of template. Figure 6 shows shaped ceramic fluid with variations in number of channels, area of patterning and dimensions of structure (see fig.6. a-c). As the number of channels/outlets can be varied, broader or narrower drug concentration gradient can be generated. Therefore, this multiscale fabricated mesh pattern based microfluidic device can also be scaled up or scaled down for concentration gradient generation.

**Figure 6:**
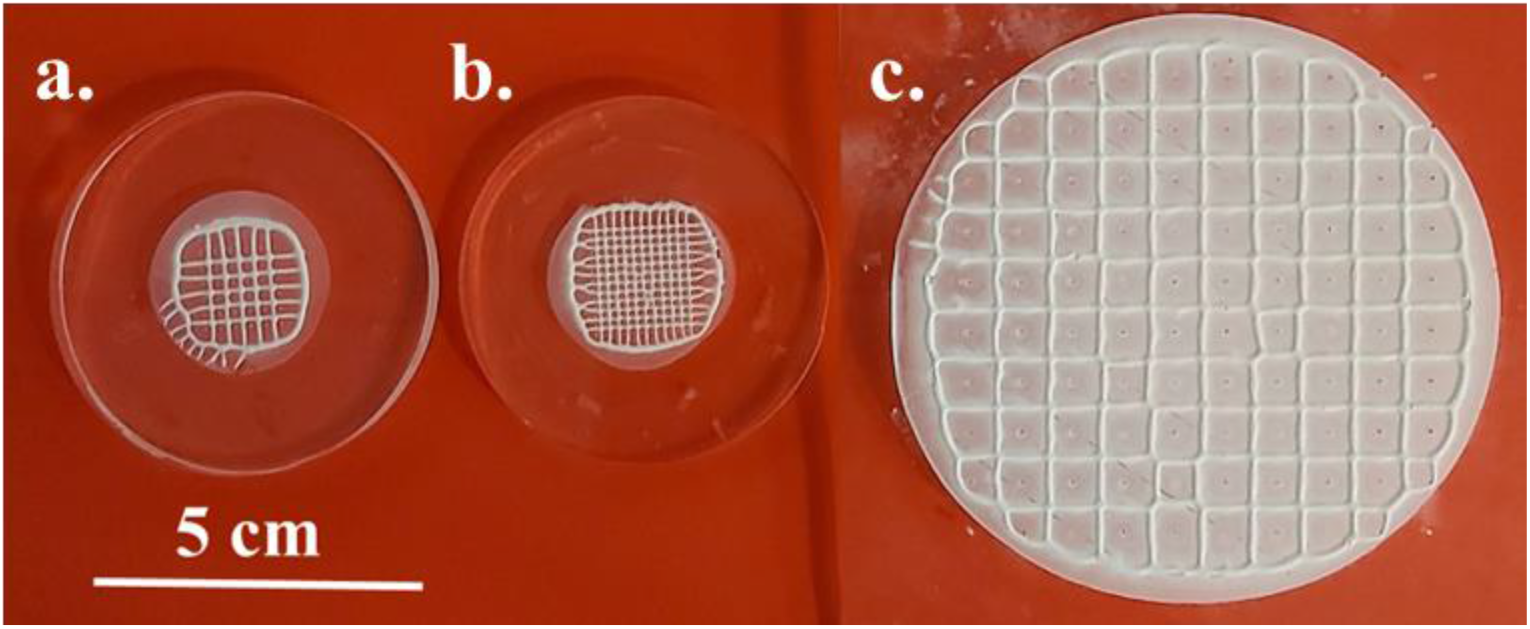
Scalability of fabrication process – (a) 5×5 mesh, (b) 11 × 11 mesh and (c) mesh patterning over large area.

## 4. Discussion

In this study, we used mesh like template generated by shaping ceramic fluid controlled via lifted Hele-Shaw for developing microfluidic concentration gradient generator (μCGG). The outlet concentration w.r.t. flow rate was determined using COMSOL simulations and validated using experiments with fluorescein dye-FITC. Next, the gradient of curcumin in growth media was estimated using the absorbance of curcumin. The drug testing application in fabricated μCGG was successfully demonstrated by measuring IC50 of curcumin on model cell line – HeLa (cervical cancer cells). Further, consistent IC50 value from both - conventional well-plate assay and microfluidic devices-depicts potential use of device in concentration dependant studies.

For generation of complex concentration gradient profile, “Christmas-tree” based designs ^19,20^ are widely used. It has been used for biological studies including cellular migration ^21^, proliferation and neuronal differentiation ^22^ and non-biological studies such as chemical synthesis ^1^, serial dilution ^2,3^, sensors ^4,5^ etc. However, most of the lithography-based templates and respective μCGG are complex and involves multi-step fabrication ^11^, thereby increasing cost and time of fabrication. To overcome these limitation, some non-conventional methods have been reported such as paper based high-throughput devices ^12^, thread based microfluidics ^14,23^ etc were reported. However, collection of solution is difficult and studies are majorly conducted inside the device. These shortcomings are addressed in the proposed microfluidic device. We have demonstrated microfluidic device with 10 outlets; however, number of channels/outlets can be easily changed by varying number and position of drilled holes, thereby controlling gradient strength and resolution. Therefore, fabrication of lithography-less template using lifted Hele-Shaw cell is advantageous in terms of scalability, cost of fabrication and customisable inlet and outlet of fabricated μCGG. This study highlights the potential of lifted Hele-Shaw cell in fabrication of microfluidic device for drug testing application and can be further explored for other biological and non-biological applications.^24^

## 5. Conclusion

Our work demonstrates that mesh-like shaping of ceramic fluids can be utilised to fabricate microfluidic devices for microfluidic concentration gradient generation. The application of fabricated μCGG was demonstrated by testing curcumin-drug gradient on cervical cancer cells (HeLa). We believe that this simple, customised number of outlets, cost effective and scalable μCGG can be used for investigating various concentration gradient-based biological and non-biological studies.

## Supporting information

The following files are available free of charge.

Figure S1 - Calibration curve of FITC intensity with respect to known FITC concentration. Figure S2 - Calibration curve of curcumin absorbance with respect to known curcumin concentration

## Author contribution

SY, KB, MR fabricated template for the study. MR did the template characterization on WLI. SY, PT performed COMSOL simulations and initial experiments. SY, KB performed drug testing experiments and analyzed data. AM and PG designed experiments, methodology and supervised the project. AM, PG, SY, PT, KB and MR contributed to the manuscript writing. The equal contributing authors can interchange the position in their respective CVs.

## Conflict of interest

The authors declare no conflict of interest.

## Acknowledgement

We acknowledge Department of Science and Technology, INSPIRE for providing fellowship to SY, Ministry of Health and Resource Development (MHRD) to MR, and DST-IMPRINT and IRCC IITB to KB. AM and PG thank Department of Science and Technology - Impacting Research Innovation and Technology (DST-IMPRINT, Project Number 6722), for providing financial support for the work presented in this paper. We also acknowledge white light interferometer facility at SML, Mechanical Engineering Department – IITB, Prof. Dulal Panda and Prof. Rahul Purwar, BSBE IITB for providing access to spectrophotometer.

## Notes

### Competing Interest Statement

The authors have declared no competing interest.

### Summary of Updates

Figure 4 and respective text revised

