## supplementary figure 1,2 for "Scalable large-area mesh-structured microfluidic gradient generator for drug testing applications"

**Abstract**

Microfluidic concentration gradient generators are useful in drug testing. drug screening, and other cellular applications to avoid manual errors, save time, and labor. However, expensive fabrication techniques make such devices prohibitively costly. Here, in the present work, we developed a microfluidic concentration gradient generator (μCGG) using a recently proposed non-conventional photolithography-less method. In this method, ceramic suspension fluid was shaped into a square mesh by controlling Saffman Taylor instability in a Multiport Lifted Hele-Shaw Cell (MLHSC). Using the shaped ceramic structure as template, the microfluidic concentration gradient generator (μCGG) was prepared by soft lithography. The concentration gradient was characterized and effect of the flow rates were studied usingCOMSOL simulations. The simulation result was further validated by creating fluorescein dye (Fluorescein isothiocanate, FITC) gradient in the fabricated μCGG. To demonstrate the use of this device for drug testing, we created various concentrations of an anticancer drug – curcumin - using the device and determined its inhibitory concentration on cervical cancer cell-line HeLa. We found that the IC50 of curcumin for HeLa to be 28.6 ± 6.1 μM which matched well with the conventional muti-well drug testing method (34.9 ± 1.7 μM). This method of μCGG fabrication has multiple advantages over conventional photolithography such as: i) the channel layout and inlet-outlet arrangements can be changed by simply wiping the ceramic fluid before it solidifies, (ii) it is cost effective, (iii) large area patterning is easily achievable, and (iv) the method is scalable. This technique can be utilised to achieve broad range of concentration gradient to be used for various biological and non-biological applications.

**Supplementary data:**

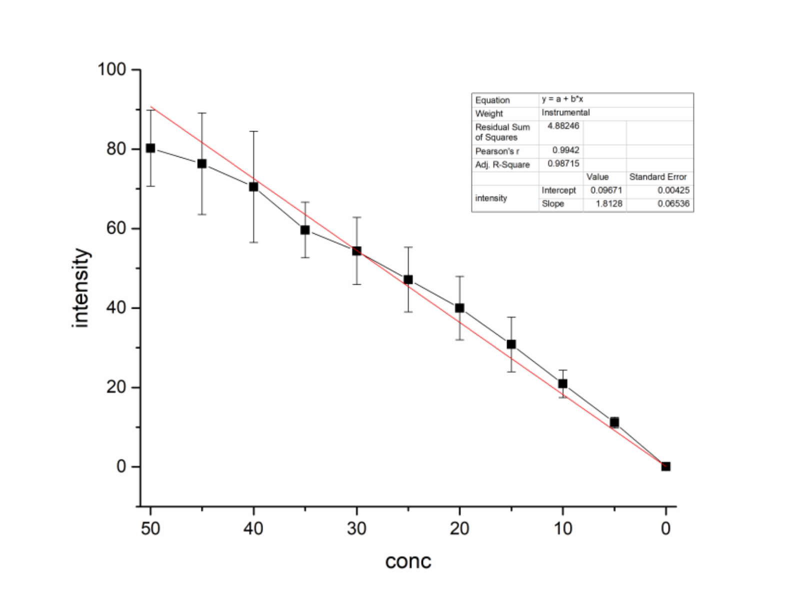


Figure S1: Calibration curve of FITC intensity with respect to known FITC concentration

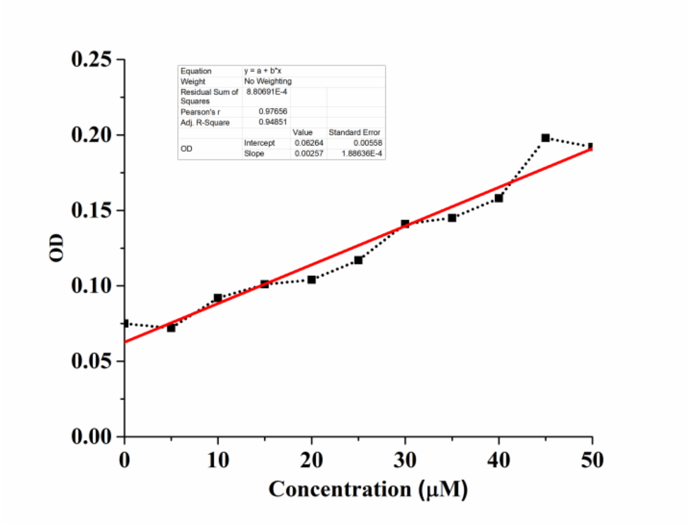


Figure S2: Calibration curve of curcumin absorbance with respect to known curcumin concentration
